# Phosphorylation of Threonine 107 by Calcium/Calmodulin dependent Kinase II δ Regulates the Detoxification Efficiency and Proteomic Integrity of Glyoxalase 1

**DOI:** 10.1101/2020.04.09.033159

**Authors:** Jakob Morgenstern, Sylvia Katz, Jutta Krebs-Haupenthal, Jessy Chen, Alireza Saadatmand, Fabiola Garcia Cortizo, Alexandra Moraru, Johanna Zemva, Marta Campos Campos, Aurelio Teleman, Johannes Backs, Peter Nawroth, Thomas Fleming

## Abstract

The glyoxalase system is a ubiquitously expressed enzyme system with narrow substrate specificity and is responsible for the detoxification of harmful methylglyoxal (MG), a spontaneous by-product of energy metabolism. Glyoxalase 1 (Glo1) is the first and therefore rate limiting enzyme of this protective system. In this study we were able to show that a phosphorylation of threonine-107 in the Glo1 protein, mediated by Ca^2+^/Calmodulin-dependent Kinase II delta (CamKIIδ), is associated with elevated catalytic efficiency of Glo1. In fact, Michaelis-Menten kinetics of Glo1 mutants revealed that a permanent phosphorylation of Glo1 was associated with increased V_max_ (1.23 µmol/min/mg) and decreased K_m_ (0.19 mM HTA), whereas the non-phosphorylatable Glo1 showed significantly lower V_max_ (0.66 µmol/min/mg) and increased K_m_ (0.31 mM HTA). This was also confirmed with human recombinant Glo1 (V_max_ (Glo1_phos_) = 999 µmol/min/mg; K_m_ (Glo1_phos_) = 0.09 mM HTA vs. V_max_ (Glo1_red_) = 497 µmol/min/mg; K_m_ (Glo1_red_) = 0.12 mM HTA). Additionally, proteasomal degradation of non-phosphorylated Glo1 via ubiquitination occurred more rapidly as compared to native Glo1. The absence of the responsible kinase CamKIIδ was associated with poor MG detoxification capacity and decreased protein content of Glo1 in a murine CamKIIδ knock-out model. Furthermore, this regulatory mechanism is also related to an altered Glo1 status in cancer, diabetes and during aging. In summary, phosphorylation of threonine-107 in the Glo1 protein by CamKIIδ is a quick and precise mechanism regulating Glo1 activity.

## Introduction

Glyoxalase 1 (Glo1) is the first enzyme of a catalytic complex described as the glyoxalase system, which is expressed in all living cells. It is mainly responsible for the detoxification of methylglyoxal (MG), a spontaneous by-product which is generated during glycolysis. MG is a highly reactive 2-oxoaldehyde and represents a precursor for advanced glycation endproducts (AGE), which are leading to increased reactive oxygen species in the cell [1]. Consequently, given the omnipresent formation of MG, the glyoxalase system represents a major mechanism in the xenobiotic metabolism to prevent oxidative stress [2].

In order to respond in an economical way to different cellular stimuli the glyoxalase system has to undergo rapid molecular adjustments. Glo1 can be nitrosylated in cooperation with glutathione, which leads to a decreased enzymatic activity; a phenomenon which has been described in crude organisms (yeast) and mammalian cells [7, 8]. Phosphorylation of Glo1 has also been found to be present in mammalian cells, yeast and in plants. In fibroblasts the phosphorylation has been linked to the induction of necrosis by TNFα [9]. In the same study threonine-107 (T107) was identified for the first time as one potential phosphorylation site, but neither the responsible kinase nor the enzymatic consequences could be shown [10]. In which way post-translational modifications of Glo1 regulate the efficiency of the glyoxalase system within various pathological contexts is currently not understood. However, alterations of Glo1 and its activity seems to play a pivotal role in various clinical contexts such as diabetes, aging, as a potential drug target regarding cancer therapeutics but also as treatment against bacteria or protozoans [3, 4]. Furthermore, psychological disorders, e.g. anxiety-like behavior as well as alcohol use disorders have been linked to altered Glo1 [5, 6]. The aim of this study was to investigate the phosphorylation of Glo1, characterize responsible kinase(s) and describe consequences *in vitro* and *in vivo*.

## Results

### Phosphorylation of Glyoxalase 1 at threonine 107 affects kinetic efficiency of methylglyoxal detoxification and proteasomal degradation rate

In order to investigate the effect of a Glo1 phosphorylation at T107, two murine cardiac endothelial cell models were established. For both cell model systems a previously established Glo1 knock-out model was used in order to prevent any endogenous Glo1 activity (supplementary Figure 1; material & methods). The permanently phosphorylated clone (P) was established with an exchange of threonine to glutamic acid T107E, whereas the non-phosphorylatable clone (NP) was established with an exchange of threonine to alanine (T107>G107) (supplementary figure 1; materials & methods). When Glo1 activity was normalized to total protein content a Michaelis-Menten kinetic revealed that a permanent phosphorylation of Glo1 was associated with increased V_max_ (1.23 µmol/min/mg) and decreased K_m_ (0.19 mM HTA), whereas the NP clone showed significantly lower V_max_ (0.66 µmol/min/mg) and increased K_m_ (0.31 mM HTA). Wild-type (WT) cells showed an enzymatic efficiency between those two clones (V_max_ = 0.95 µmol/min/mg; K_m_ = 0.24 mM HTA) reflecting potentially an intermediate state of Glo1 phosphorylation (Figure 1 A). Regarding intracellular MG concentrations NP clones showed an approximately 50% increase as compared to the WT- and P-clones (Figure 1 B). Using flow cytometry and dichlorofluorescein, the intracellular ROS levels were quantified, which revealed that NP-clones have significantly higher ROS levels (151 ± 13 a.u.) as compared to P-clones (108 ± 11 a.u.) and WT cells (100 ± 7 a.u.) (Figure 1 C). As a consequence NP-clones showed nuclear damage, displayed by significantly increased tail length in a comet assay (Figure 1 D) and increased p53- as well as γH2Ax- expression in cells lacking Glo1 phosphorylation (NP) (Figure 1 E). This resulted also in lower proliferation rates as measured by bromo deoxyuridine incorporation, in which NP clones displayed only ∼58% proliferation rate of the WT cells (Figure 1 F).

**Figure 1.**
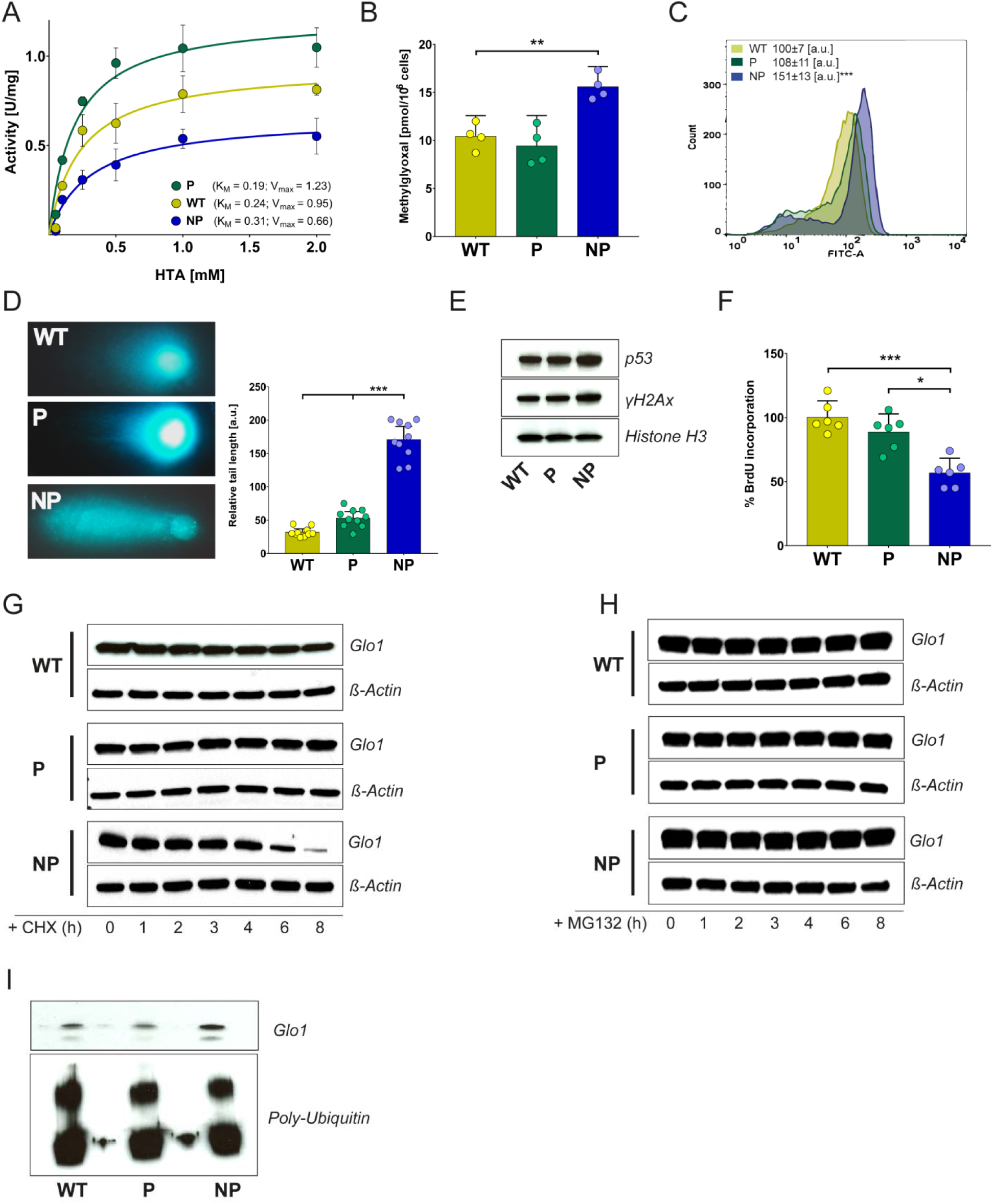
Phosphorylation of Glyoxalase 1 at threonine 107 affects kinetic efficiency of methylglyoxal detoxification. **A**, kinetic profile of the Glo1 catalysed reduction of hemithioacetal in wild-type (WT), phosphorylated (P) and non-phosphorylated (NP) clones. **B**, intracellular MG-levels in wild-type (WT), phosphorylated (P) and non-phosphorylated (NP) clones cultured under baseline conditions (5 mM Glucose). **C**, intracellular levels of reactive oxygen species in wild-type (WT), phosphorylated (P) and non-phosphorylated (NP) clones under baseline conditions (5 mM Glucose) using flow cytometry and H_2_DCFDA as reagent. **D**, left: Oxidation of cellular DNA measured by Comet Assay in wild-type (WT), phosphorylated (P) and non-phosphorylated (NP) clones under baseline conditions (5 mM Glucose). Right: Relative tail length of appropriate comets (n=15) in wild-type (WT), phosphorylated (P) and non-phosphorylated (NP) clones under baseline conditions (5 mM Glucose). **E**, representative western blot analysis of total cell extracts (30 µg of protein) from wild-type (WT), phosphorylated (P), non-phosphorylated (NP) clones and from wild-type cells probed with anti-p53 antibody, anti-γH2Ax antibody and anti-β-Actin antibody as a loading control. **F**, median proliferation rate in wild-type (WT), phosphorylated (P) and non-phosphorylated (NP) clones under baseline conditions (5 mM Glucose). **G**, representative western blot analysis of total cell extracts (30 µg of protein) from wild-type (WT), phosphorylated (P) and non-phosphorylated (NP) clones after cycloheximide (CHX; 10µg/mL) treatment probed with anti-Glo1antibody and anti-ß-Actin antibody as a loading control. **H**, representative western blot analysis of total cell extracts (30 µg of protein) from wild-type (WT), phosphorylated (P) and non-phosphorylated (NP) clones after MG-132 treatment (10 µM) probed with anti-Glo1antibody and anti-ß-Actin antibody as a loading control. **I**, representative western blot analysis of total cellextracts (100 µg of protein) from wild-type (WT), phosphorylated (P) and non-phosphorylated (NP) clones after an ubiquitin-pull-down approach probed with anti-Glo1antibody and anti-Poly-Ubiquitin antibody as a loading control. All data represent the mean of 4-10 independent experiments ± standard deviation. *** p < 0.001; ** p < 0.01; * p < 0.05.

In order to investigate whether Glo1 protein stability was affected by the phosphorylation status of T107, cycloheximide (CHX), a protein synthesis inhibitor, was used. During the treatment NP-clones showed a rapid degradation of Glo1 protein as compared to the WT clones; whereas in P-clones Glo1 protein was not changed after 8 hrs of CHX treatment (Figure 1 G). Using a proteasome inhibitor (MG132), NP-clone showed no change in protein content comparable to WT and P-clones after 8 hrs of treatment (Figure 1 H). To confirm that it is the rapid degradation of Glo1 via ubiquitinylation in the NP-clones, a ubiquitin-pull-down experiment revealed that NP-clones had a significantly higher concentration of Glo1 in the ubiquitin isolated fraction (Figure 1 I).

### Phosphorylation of Glyoxalase 1 is mediated by CamKIIδ in vitro and in vivo

Preliminary results (data not shown) suggested that CamKII is a suitable candidate in order to investigate the effect of a Glo1 phosphorylation. A [γ-^32^P]-ATP Kinase assay revealed that Ca^2+^/calmodulin-dependent protein kinase II δ (CamKIIδ) can phosphorylate recombinant human Glo1 protein in a dose-dependent matter (Figure 2 A). Using recombinant human Glo1, a Michaelis-Menten kinetic was performed in order to compare the results with kinetics obtained from Glo1 mutants (see Figure 1). In line with the previous results it showed a two-fold increased V_max_ and an increased affinity of phosphorylated Glo1 in comparison to reduced Glo1 (V_max_ (Glo1_phos_) = 999 µmol/min/mg; K_m_ (Glo1_phos_) = 0.09 mM HTA vs. V_max_ (Glo1_red_) = 497 µmol/min/mg; K_m_ (Glo1_red_) = 0.12 mM HTA) (Figure 2 B). The pharmacological inhibition of CamKII via KN93 in endothelial cells showed a decline in Glo1 activity and protein content after 24 hrs (Figure 2 C & E). This was accompanied by a mild, but significant, increase (∼20%) in intracellular MG concentrations (Figure 2 D). An overexpression of CamKIIδ was not linked to an increase of Glo1 activity (Figure 2 F). Using a Phos-tag approach, a major shift in Glo1 band was observed in recombinant human Glo1 incubated with CamKIIδ and ATP as well as cells treated with KN93. Validity of this band-shift caused by altered phosphorylation status was confirmed using λ-Protein Metallo-Phosphatase where the upper band disappeared (Figure 2 G).

**Figure 2.**
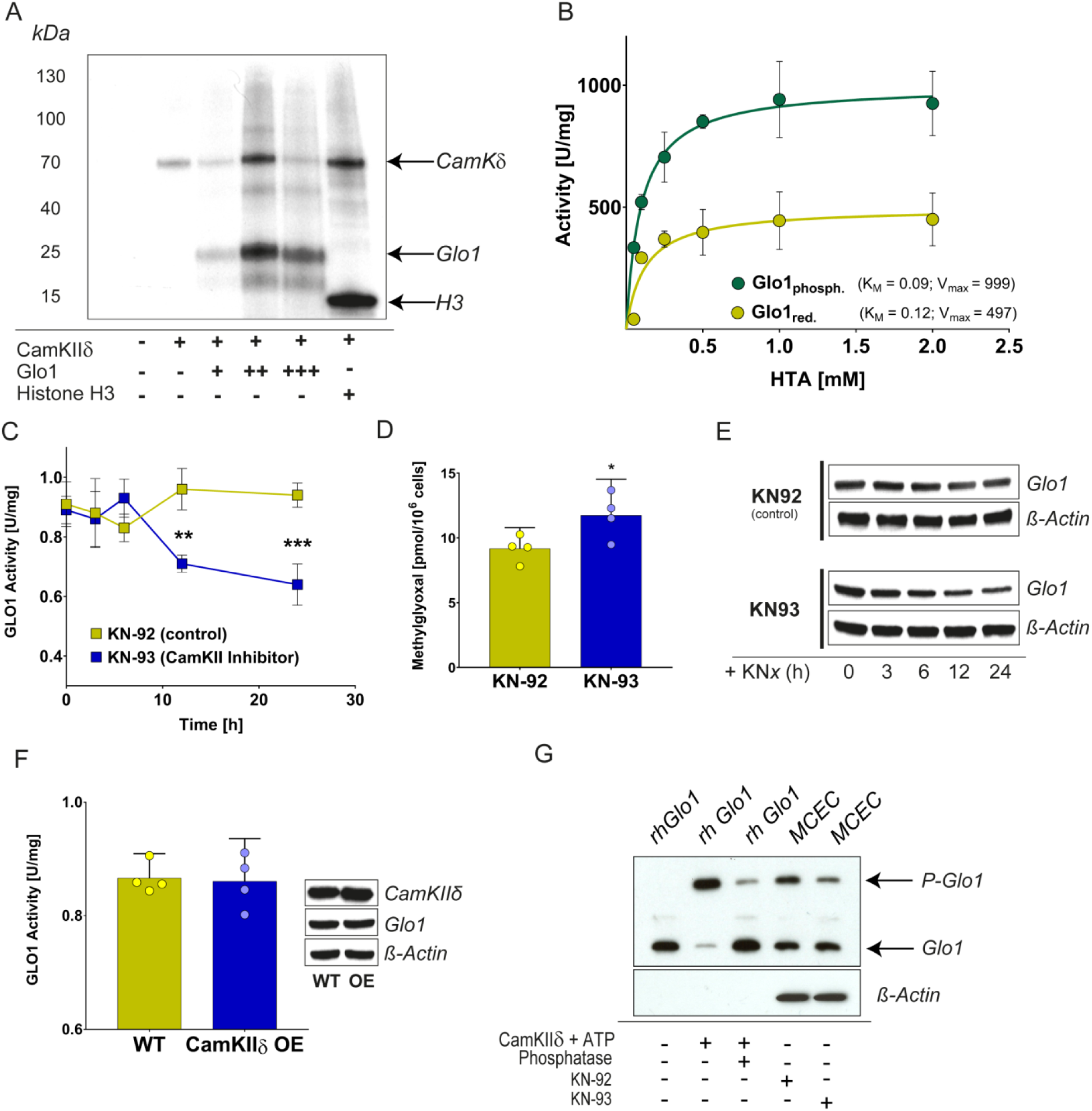
Phosphorylation of Glyoxalase 1 is mediated by CamKIIδ *in vitro* and *in vivo*. **A**, representative autoradiography blot of recombinant human Glo1 protein using radioactive ATP (γ-^32^P) with & without CamKIIδ and Histone H3 as control. **B**, kinetic profile of the Glo1 catalysed reduction of hemithioacetal using phosphorylated recombinant human protein (Glo1_phos_) and unphosphorylated recombinant human protein (Glo1_red_). **C**, Glo1 catalysed reduction of hemitioacetal in wild-type cells 6-24 hrs after specific KNx treatment. **D**,. intracellular MG-levels in wild-type cells after specific KNx treatment. **E**, representative western blot analysis of total cell extracts (30 µg of protein) from wild-type cells after specific KNx treatment probed with anti-Glo1antibody and anti-ß-Actin antibody as a loading control. **F**, Glo1 catalysed reduction of hemitioacetal in wild-type cells 12 hrs after over-expression (OE) of CamKIIδ. **G**, representative western blot analysis of cytosolic cell extracts (30 µg of protein) using a Phos-Tag-Gel (Zinc) approach of recombinant human Glo1 and wild-type cells (MCEC) after specific KNx treatment probed with anti-Glo1 antibody and anti-ß-Actin antibody as a loading control. All data represent the mean of at least 4 independent experiments ± standard deviation. *** p < 0.001; ** p < 0.01; * p < 0.05

### CamKIIδ knock-out model reflects a loss of Glo1 protein/activity due to missing phosphorylation status

A murine model with a global CamKIIδ knock-out (KO) was used to investigate the impact of a total absence of CamKIIδ towards Glo1 protein and its phosphorylation status. In 20-weeks old male C57BL/6 CamKIIδ KO mice, the protein content of Glo1 was globally reduced by approximately 50% compared to control animals, with the liver and heart being the most affected (Figure 3 A). This was also confirmed by Glo1 enzyme activities in those organs (Figure 3 B). The observed downregulation of Glo1 protein and activity in CamKIIδ KO mice was associated with a decrease in its phosphorylation status in the liver tissue (Figure 3 F). Interestingly, the CamKIIδ KO mice model was neither linked to increased MG nor MG-H1 concentrations in whole tissue lysates (Figure 3 C & D). However, potential nuclear damage was shown by increased p53-, but not γH2Ax-expression in those tissues as compared to *in vitro* results (Figure 3 E).

**Figure 3.**
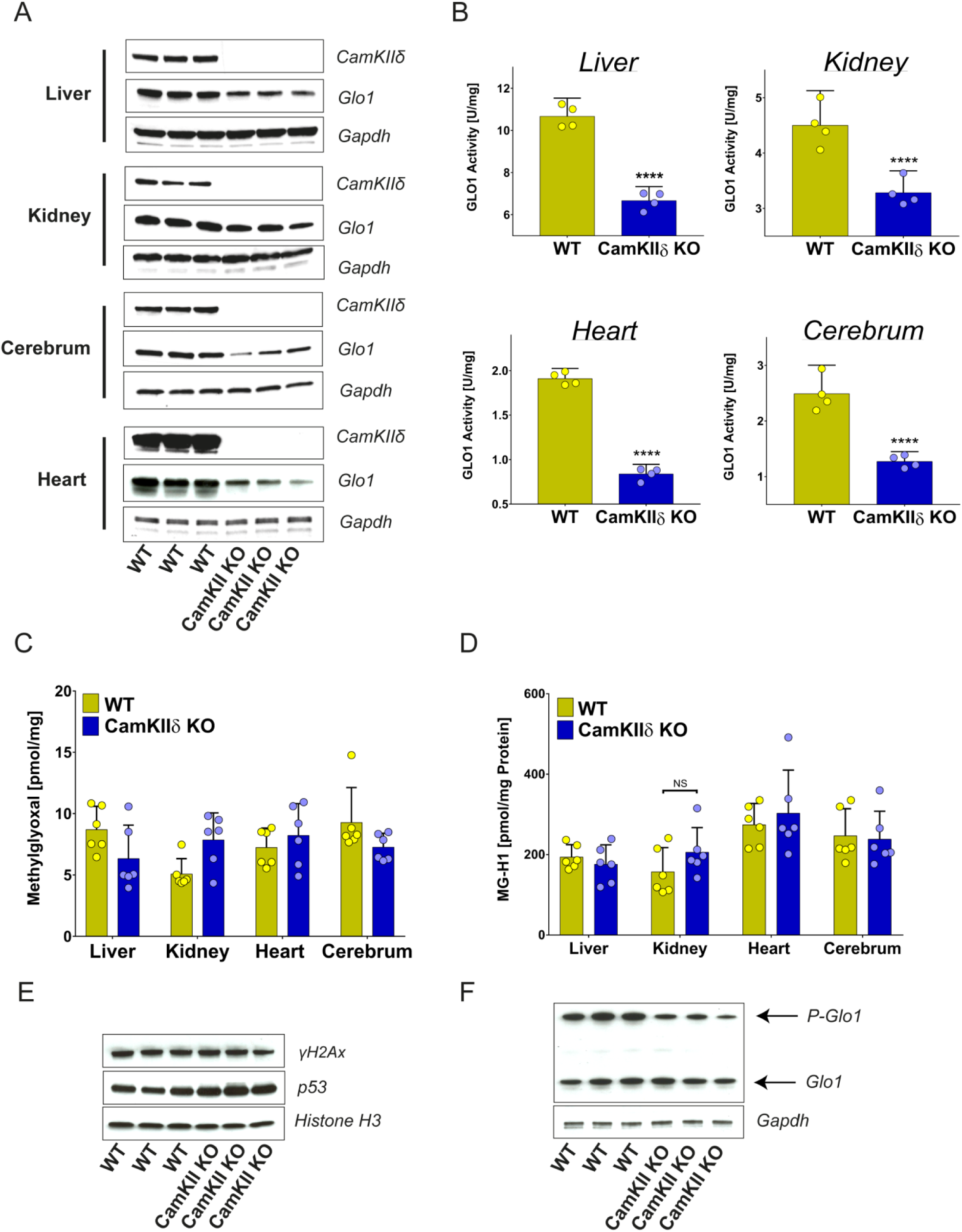
Unphosphorylated Glo1 is linked to nuclear MG accumulation and damage. **A**, representative western blot analysis of cytosolic cell extracts (30 µg of protein) of various tissues from wild-type (WT) and CamKIIδ KO mice probed with anti-CamKIIδ antibody, anti-Glo1 antibody and anti-GAPDH antibody as a loading control. **B**, Glo1 catalysed reduction of hemitioacetal in cytosolic cell extracts of various tissues from wild-type (WT) and CamKIIδ KO mice. **C**, MG levels in various whole tissue sections from wild-type (WT) and CamKIIδ KO mice. **D**, MG-H1 levels in various whole tissue sections from wild-type (WT) and CamKIIδ KO mice. **E**, representative western blot analysis of total cell extracts (30 µg of protein) of liver tissue from wild-type (WT) and CamKIIδ KO mice probed with anti-p53 antibody and anti-Histone H3 antibody as a loading control. **F**, representative western blot analysis of cytosolic liver extracts (30 µg of protein) using a Phos-Tag-Gel (Zinc) approach of wild-type (WT) and CamKIIδ KO mice probed with anti-Glo1 antibody and anti-GAPDH antibody as a loading control. All data represent the mean of 4-6 independent experiments ± standard deviation. **** p < 0.0001

### Glo1 activity and protein status is altered in diabetes, cancer or during aging and is linked to its phosphorylation status

In type 1 (Streptozotocin (STZ)) and type 2 (leptin deficient (db/db) diabetic mouse model it was revealed that liver tissue showed significantly decreased Glo1 activity as compared to the WT controls already at 20 weeks of age (Figure 4 A & B). This decline was most pronounced in liver tissue, but was a global phenomenon observable in all major tissues (data not shown). In both diabetic animal models the decrease in Glo1-activity and protein content in the liver was also associated with a lower phosphorylation status of Glo1 as compared to WT animals (Figure 4 D & E). Furthermore, Glo1 activity also declined during aging with approximately 50% reduction in 80 weeks old WT mice as compared to 10 weeks old wild-type mice (Figure 4 B). Again, this phenomenon was linked to a lack of Glo1 phosphorylation during aging (Figure 4 E). In addition to ageing and diabetes, altered Glo1 activity has been described frequently in various malignant solid tumors. In two human cell lines derived from breast cancer (MCF-7) and cervical cancer (HeLa) Glo1 activity and protein content were screened. The comparison with human umbilical endothelial cells (HUVECs) revealed a 2.5- (MCF-7) and 1.4-(HeLa) fold higher Glo1-activity and Glo1 protein content of the cancerous cell lines (Figure 4 C & D). Especially in MCF-7 cells it was shown an increased state of Glo1 phosphorylation as compared to HUVECS, whereas HeLa cells showed an intermediate state of Glo1 phosphorylation (Figure 4 E). Regarding the expression status of CamKIIδ we found reduced mRNA levels in liver tissue of both diabetic animal models, but a highly increased CamKIIδ expression in cancerous cell lines as compared to HUVECs (Figure 4 F).

**Figure 4.**
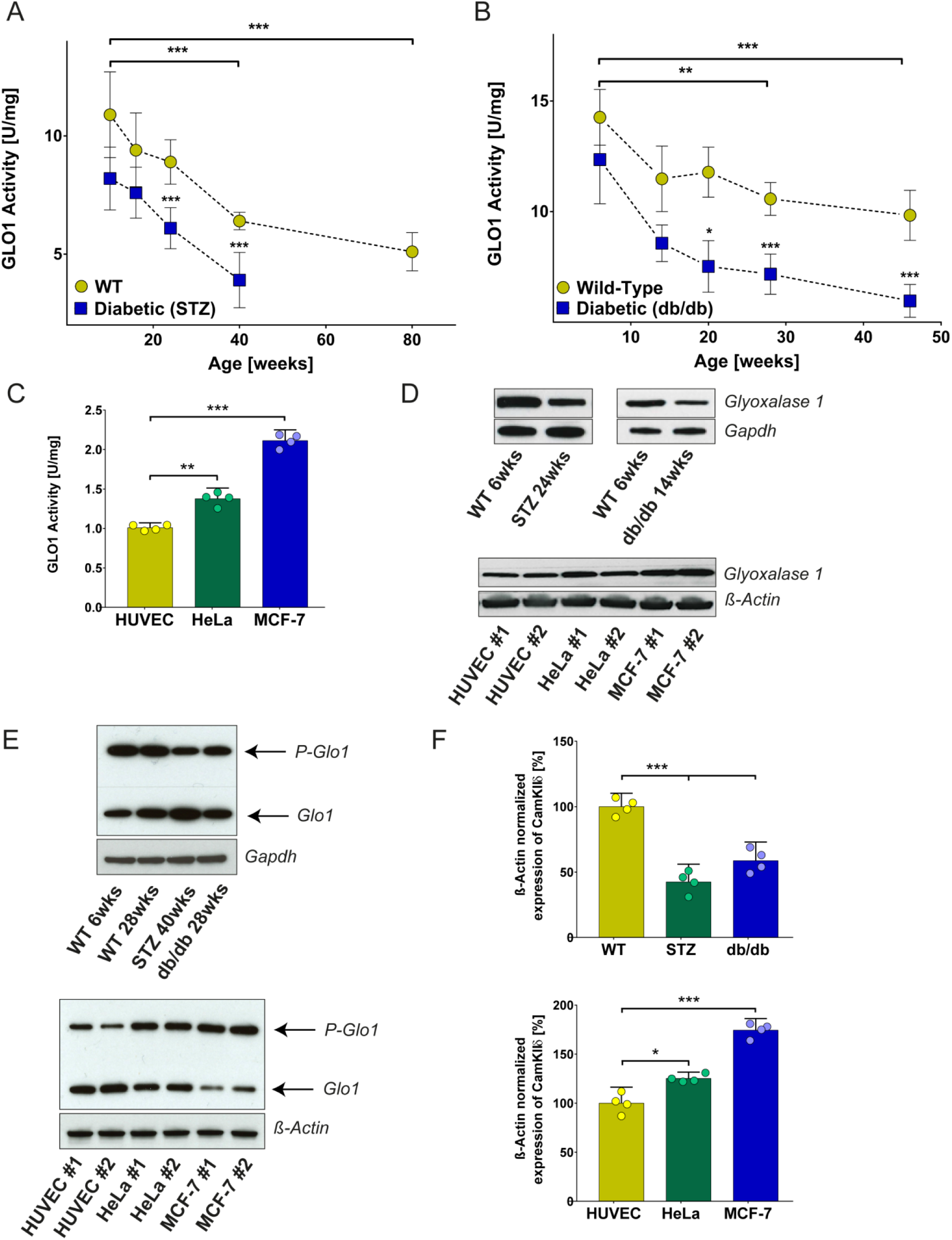
Glo1 activity and protein status is altered in diabetes, cancer and during aging and is linked to its phosphorylation status. **A**, Glo1 activity in liver tissue of wild-type (WT) and diabetic (STZ) mice between 10 and 80 weeks of age. **B**, Glo1 activity in liver tissue of wild-type (WT) and diabetic (db/db) mice between 6 and 46 weeks of age. **C**, Glo1 catalysed reduction of hemithioacetal in Human Umbilical Vein Endothelial Cells (HUVEC), cervical cancer cells (HeLa) and breast carcinoma endothelial cells (MCF-7). **D**, top; representative western blot analysis of cytosolic cell extracts (30 µg of protein) from liver tissue of wild-type mice (WT), STZ treated mice and leptin deficient mice (db/db) of different age (e.g. 6 wks - 6 weeks of age) probed with anti-Glo1 antibody and anti-GAPDH antibody as a loading control. Bottom; representative western blot analysis of cytosolic cell extracts (30 µg of protein) of Human Umbilical Vein Endothelial Cells (HUVEC), cervical cancer cells (HeLa) and breast carcinoma endothelial cells (MCF-7) probed with anti-Glo1 antibody and anti-ß-Actin antibody as a loading control. **E**, top; representative western blot analysis of cytosolic liver extracts (30 µg of protein) using a Phos-Tag-Gel (Zinc) approach of wild-type mice (WT), STZ treated mice and leptin deficient mice (db/db) of different age (e.g. 6 wks - 6 weeks of age) probed with anti-Glo1 antibody and anti-GAPDH antibody as a loading control. Bottom; representative western blot analysis of cytosolic liver extracts (30 µg of protein) using a Phos-Tag-Gel (Zinc) approach of Human Umbilical Vein Endothelial Cells (HUVEC), cervical cancer cells (HeLa) and breast carcinoma endothelial cells (MCF-7) probed with anti-Glo1 antibody and anti-ß-Actin antibody as a loading control. **F**, top; mRNA expression of CamKIIδ in liver tissue of wild-type (WT), type 1 diabetes (STZ) and type 2 diabetes (db/db) mice normalized to ß-Actin. Bottom; mRNA expression of CamKIIδ in Human Umbilical Vein Endothelial Cells (HUVEC), cervical cancer cells (HeLa) and breast carcinoma endothelial cells (MCF-7) normalized to ß-Actin. All data represent the mean of at least 4 independent experiments ± standard deviation. *** p < 0.001; ** p < 0.01; * p < 0.05

**Figure 5.**
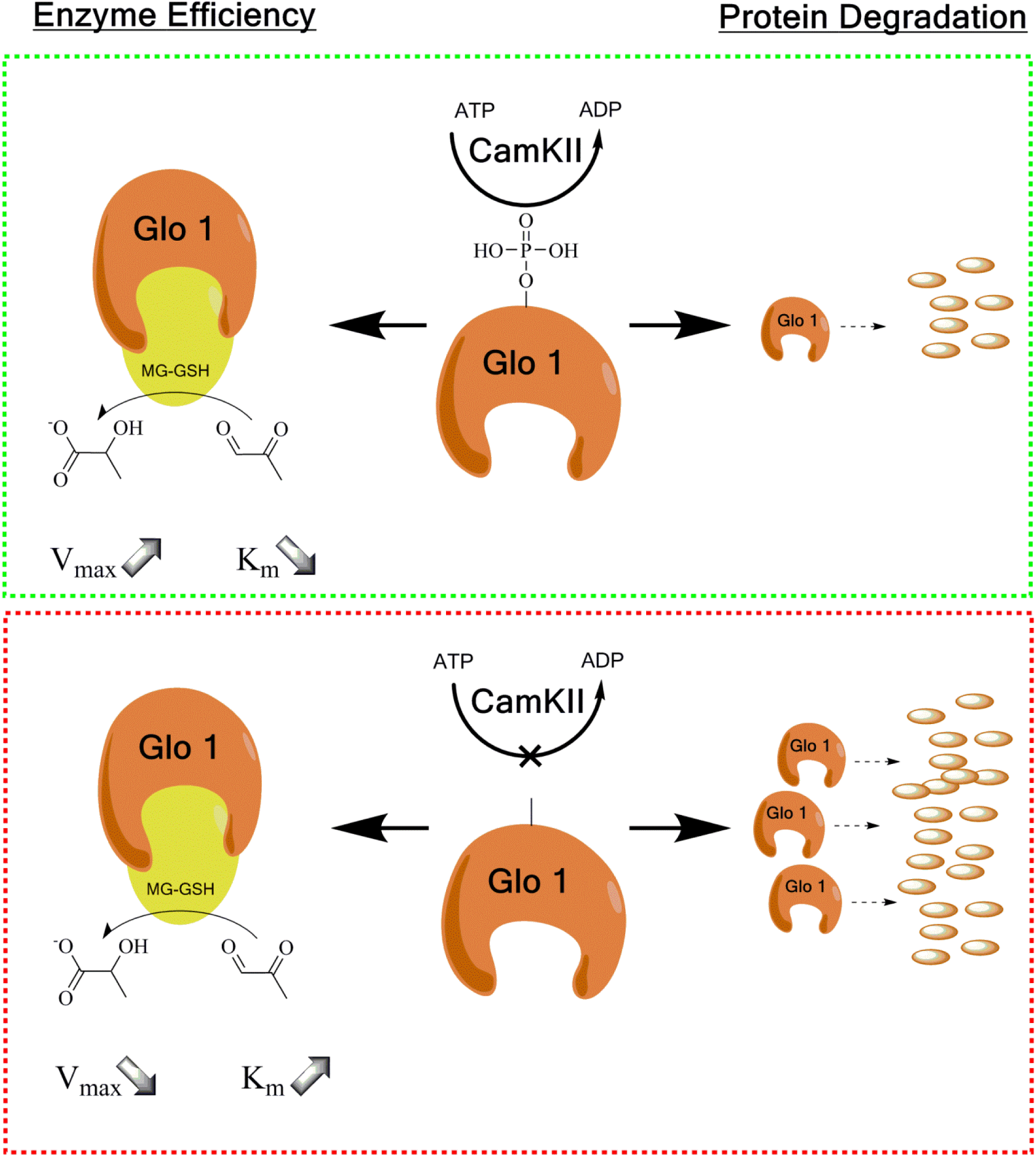
Proposed mechanism of action

## Discussion

Due to its ubiquitous existence in all living cells (Glo1 is in the top 10% of intracellular protein abundancy), it is believed that the glyoxalase system has a highly conserved and therefore fundamental role [16, 17, 20]. The associations between the glyoxalase system, in particular Glo1, and patho-mechanisms in diabetes, cancer, anxiety, aging, HIV or drug abuse are striking and suggest that Glo1 has a central role in the maintenance of molecular homeostasis [6, 8, 9, 10]. However, after decades of research it seems the only conclusive function of the glyoxalase system is the detoxification of MG, a by-product of energy metabolism [7]. In addition, independent studies showed in various Glo1 KO models that this enzyme system is not indispensable for living cells due to effective compensatory detoxification of harmful MG [14, 15, 19, 21].

Post-translational modifications of Glo1, have generally been overlooked as a regulator of its biological function, with only a few limited studies have been conducted. In plants (*arabidopsis thaliana*) Glo1 was phosphorylated via the kinase SnRK2.8, a kinase involved in stress response. This phosphorylation of Glo1 was associated with an increase in enzyme activity [16]. The treatment of yeast with mating factor resulted in the same consequences; a phosphorylated Glo1 with increased enzymatic activity [17]. In mammals the situation seems to be more complex and experimental results are rare, as well as partly inconsistent. In a redox-dependent regulation it was found that Glo1 can be glutathionylated, but is dependent on the addition of NO, which consequently leads to the formation of S-nitrosoglutathione through Glo1. The glutathionylation of Glo1 at cysteine 139 and the formation of S-nitrosoglutathione are both leading to an inhibition of enzymatic activity [12, 18]. Subsequent studies were able to identify a phosphorylation site of Glo1 in L929 cells; a necessity for the induction of necrosis by tumor necrosis factor alpha [9, 10]. Within this context, the authors of this study speculated that phosphorylation of Glo1 is necessary for cell death and that this is either driven by protein kinase A, glycogen synthase kinase 3 or CamKII [9, 11]. Interestingly, this study did not find any evidence for altered enzyme kinetics driven by phosphorylation. Methodical limitations (assay was performed under substrate saturation) could be an explanation for this contradictory finding. Furthermore, the authors’ hypothesis “Glo1 inducing cell death” can be interpreted as controversial, given the background that Glo1 has a highly conserved protective function within the cell.

The experimental study herein presents evidence that CamKIIδ is a major regulator of the Glo1 phosphorylation status. This phenomenon has significant consequences for enzymatic efficiency and proteasomal degradation of Glo1 *in vivo*. In fact phosphorylation of threonine 107 leads to an optimized enzymatic efficiency with higher V_max_ and lower K_m_ values. From a stereochemical viewpoint, a strong negative charge from a phosphate group alters how a given protein is shaped, but more important how it interacts with water. When an enzyme becomes more hydrophilic it can interact with hydrophilic substrates more efficiently. The substrate of Glo1, MG, occurs under physiological conditions >99% in a mono- and dihydrated form, which binds it easily to the cysteine group of GSH [22]. Additionally, the catalytic mechanism suggests that the activity of Glo1 is driven by the capacity of the hydrogen bonding network of glutamate 172, which is likely the proton abstracting base within the catalytic mechanism. In line with recent stoichiometrically findings, it can be hypothesized that the hydrophilic environment due to phosphorylation of threonine 107 points towards optimized kinetic conditions for the removal of a proton from carbon 1 of the hemithioacetal moiety of the substrate and consecutive transfer to carbon 2 of the same molecule [23].

Moreover, the experiments herein showed that decreased Glo1 phosphorylation is highly associated with declined Glo1 activity in aging or diabetes and that the opposite effect seems to take place in human tumor cells, where Glo1 activity is highly upregulated. In line with that, CamKIIδ expression shows also an increase in tumor cells and a downregulation in diabetic animal models. Therefore, we provide for the first time an explanation for phenomena, which have been described for a long time in a broad range of experimental and clinical contexts [4, 5, 6, 24, 25, 26, 27]. Further studies should focus on CamKIIδ expression and the effect towards activity of Glo1 in order to reveal new molecular linkages.

## Materials & Methods

### Cell culture

Human HeLa cells derived from cervical cancer, MCF-7 cells derived from a human breast cancer and primary murine cardiac endothelial cells (MCEC) immortalized with SV40 large T antigen were obtained from ATCC®. Primary human umbilical vein endothelial cells (HUVECs) were isolated from the vein of the umbilical cord of pooled donors (PromoCell®). Cells were grown in DMEM (gibco) with 1 g/mL glucose (MCEC) or 4.5 g/mL (HUVEC, HeLa, MCF-7) containing 10% FCS (Sigma), 1% penicillin (10000 Units/ml) (gibco), 1% streptomycin (10 mg/ml) (gibco), 1% amphotericin B (250 µg/ml) (gibco) and 1 mM HEPES (gibco) at 37°C in a saturated humidity atmosphere containing 95% air and 5% CO_2_. Cells were grown to 60% confluence for *in vitro* experiments and passaged at 80% confluence using 0.05% Trypsin-EDTA (gibco) for a maximum of four consecutive passages.

### Generation of non-phosphorylatable (NP) and permanent phosphorylated (P) Glo1 mutants

Murine cardiac endothelial cells (MCEC) with a complete Glo1 knock-out (KO) were established using CRISPR/Cas9 technique as described previously [19]. Amino acid substitution in Glo1 KO MCECs was then achieved permanently by site-directed mutagenesis carried out by Eurofins Scientific®. Briefly, Glo1 KO MCECs were transfected with WT plasmids for Glo1, whereas NP-mutants were transfected with a plasmid leading to an exchange of threonine to glycine (T107G); P-mutants were transfected with a plasmid leading to an exchange of threonine to glutamic acid (T107E). Cell clones were isolated by single cell seeding in a serial dilution approach. After at least 5 passages mutant colonies were picked and screened for Glo1 activity and protein.

### Preparation of total/cytosolic protein extracts

For cytosolic extracts 500 μl cold lysis-buffer (10 mM HEPES, 1.5 mM MgCl2, 10 mM KCl, 0.5 mM DTT, 0.05% NP40 supplemented with a premade protease/phosphatase inhibitor cocktail (Sigma) including AEBSF, Aprotinin, Bestatin, E-64, Leupeptin, Pepstatin A) was added to 3×10^6^ cells or 30 µg of pulverized tissue. Samples were then homogenized by passing the lysate 20 times through a 20G needle. After centrifugation (8000 rpm, 10 min, 4°C) the supernatant was used for protein determination and further analysis. For total extracts 500 μl of cold Radioimmunoprecipitation buffer (RIPA; 50 mM Tris-HCl; pH 7.5, 150 mM NaCl, 1% NP40, 0.5% sodium deoxycholate, 0.1% SDS, 0.5 mM DTT, 1000 units benzonase) was used and supplemented with a premade protease inhibitor cocktail (see above). 4×10^6^ cells or 30 µg of pulverized tissue were vortexed and sonicated for 30 seconds (50% power, 3 cycles) with an ultrasonic homogenizer HD2070 (Bandelin). After 30 min of incubation samples were centrifuged (14000 rpm, 10 min, 4°C) and supernatant was used for protein determination and further analysis. All protein concentrations were determined using the Bradford technique and BSA as calibration standard as described previously [28].

### Glo1 activity assay

Activity of GLO1 was determined spectrophotometrically as described previously [29]. Briefly, the method monitors the initial rate of change in absorbance at 235 nm caused by the formation of S-D-lactoylglutathione through catalysis of Glo1. For Michaelis-Menten kinetics the assay mixture contained 0.1 - 2 mM MG (enzyme activity only with substrate saturating conditions; 2 mM MG) and 2 mM GSH in sodium phosphate buffer (50 mM, pH 6.6, 37°C) and was incubated for 15 min in advance to guarantee the complete formation of hemithioacetal. After the addition of the cytosolic protein fraction (1 μg/μl) the change in absorbance was monitored for 15 min. The activity of Glo1 described in units (U), where 1 U is the amount of GLO1 which catalyzes the formation of 1 μmol of S-D-lactoylglutathione per minute. Recombinant human Glo1 (ab87413) and recombinant human CamKIIδ (ab84552) for Glo1 kinetic analysis were purchased from Abcam.

### Quantification of methylglyoxal (MG) and methylglyoxal-derived hydroimidazolone (MG-H1)

The quantification of MG and MG-H1 by stable isotopic dilution analysis via LC-MS/MS was described previously [30, 31].

### Quantification of reactive oxygen species

Determination was based upon analysis via flow cytometry/FACS. All incubation and washing steps of living cells were done in Krebs Ringer HEPES buffer (KRH) including 136 mM NaCl, 4.7mM KCl, 1.25mM CaCl2, 1.25mM MgSO4, 10mM HEPES, 0.1% Fatty Acid Free BSA; pH7.4. Cells were stained with Hoechst 33258 NucBlue® (Thermo) for the detection of viable cells. Determination of reactive oxygen species was achieved incubating MCECs with CM-H2DCFDA (5 μM in KRH buffer) for 30 min under reduced light conditions. After 2 washing steps, cells were trypsinized and resuspended in 1 ml FACS-Buffer (10% FCS, 1 mM EDTA in PBS). Analysis of fluorophores was performed using a LSRII flow cytometer (BD Biosciences) by gating initial cell population via forward scatter against side scatter signals and detecting viable cells (Hoechst positive) by violet laser (Excitation: 405 nm; Filter: 450/40 nm). Hoechst positive cells were then analyzed for CM-H2DCFDA by a blue laser (Excitation: 488 nm; Filter: 530/30 nm).

### Determination/Visualization of DNA damage

The determination and visualization of DNA damage was achieved using the comet assay as described previously [32].

### Western blotting

20 μg protein was incubated in 5x Laemmli buffer (Sigma) at 95°C for 10 min and separated by a Mini-PROTEAN® TGX (Bio-Rad) precasted gel (4-20% acrylamide). Proteins were then transferred to a nitrocellulose membrane and blocked with 2% dry milk (in PBS) at room temperature for 1 h. Membranes were then incubated overnight at 4°C with antibodies against Glo1 (1:1000 dilution; ab137098; rabbit; Abcam), p53 (1:1000 dilution; ab131442; rabbit; Abcam), CamKIIδ (1:1000 dilution; ab181052; rabbit; Abcam), γH2aX (1:1000 dilution; 9718S; rabbit; Cell Signaling Technology), Histone H3 (1:2500 dilution; 4499S; rabbit; Cell Signaling Technology), Gapdh (1:2500 dilution; 5174S; rabbit; Cell Signaling Technology) Beta-actin (1:2500 dilution; 4967S; rabbit; Cell Signaling Technology) in 2% dry milk containing PBS and 0.05% Tween20 (PBS-T). After 3 washing steps (5 min each) with PBS-T membranes were incubated with horseradish-linked goat anti-rat (1:2000 dilution; 7077S; Cell Signaling Technology) or goat anti-rabbit (1:2000 dilution; 7077S; Cell Signaling Technology) antibody for 1h at room temperature. Proteins were visualized on X-Ray films using ECL detection reagents (GE healthcare) with varying exposure time (0.1 – 2 min).

### Separation and detection of phosphorylated Glo1

Due to a lack of specificity, we were not able to produce a monoclonal phopsho-specific antibody against murine or human Glo1, even after commercial approaches (Monoclonal Antibody Core Facility, Helmholtz Zentrum Munich). The use of Phospho-tag gels was established, where a phosphorylatable protein shows a significant shift in the gel due to the included tags. Therefore, separation and detection of phosphorylated Glo1 was carried out as described previously with minor changes [33]. Briefly, we casted 10% SDS PAGE-Gels and instead of manganese (Mn^2+^-Phos-tag) we used zinc (Zn^2+^-Phos-tag) which resulted in a better resolution of the proteins in the gel and better reproducibility after transfer.

### In vitro proliferation rate

Determination of *in vitro* proliferation rate was achieved using 5-bromodeoxyuridine (BrdU) incorporation as described previously [34].

### In vitro Calcium/Calmodulin dependent Kinase II δ (CamKIIδ) assay

Kinase assays for CamKIIδ have been performed as described previously with minor changes [35]. Briefly, CamKIIδ kinase assays were performed in 20 μl reaction volume with 1× kinase buffer (0.5 mM MOPS, pH 7, 0.1% BSA, 1 µM Calmodulin, 1 mM CaCl_2_, 10 mM MgCl_2_, 100 µM [γ-32P]ATP (∼1 Ci/mmole)). In each reaction tube, WT or human recombinant Glo1 was included as substrates (1 μg/reaction) and 1 ng of CamKIIδ kinase. The kinase reaction was conducted at 30°C for 10 min and stopped by adding equal volume of urea solution (6 M) and the unincorporated label was removed by TCA precipitation of the proteins. The pellet was washed in ice cold acetone and dried. The pellet was then re-suspended in 20 μl of 1× PBS and processed with Laemmli’s buffer. The kinase assay products were separated on 12% SDS-PAGE. The dried gel was used for Autoradiography.

### Isolation of Ubiquitin

Isolation of polyubiquitin protein conjugates was achieved using a commercial Pierce– Ubiquitin Enrichment Kit (Thermo) according to the manufacturer’s instruction. Isolated total protein fractions (∼30µg) were then used for western blotting and an anti-ubiquitin antibody (rabbit; included in Kit) was used as a loading control.

### Overexpression of CamKIIδ

Host *E.coli* strain DH10B including a mammalian expression vector (pCMV-SPORT6) for CamKIIδ (Horizon Discovery; Clone ID: MMM1013-202706167; Insert Sequence: BC042895) was plated out on LB-plates including 100 μg/ml ampicillin and incubated overnight at 37°C. Three individual clones were picked and amplified in LB broth including antibiotics for 12 h at 37°C and isolated using GenElute– HP Plasmid MaxiPrep Kit (Merck). Integrity of purified expression constructs was validated by gel electrophoresis and the concentration was determined by absorbance measurement. 1×10^6^ murine cardiac endothelial cells were prepared for transfection using a NEON® electroporation transfection system (Thermo) with the following conditions; pulse voltage: 1,300 mV, pulse width: 20 ms, pulse number: 2. Schwann cells were either transfected with an empty plasmid containing only a sham vector (wild-type) or with the plasmid containing CamKIIδ (CamKIIδ OE).

### Quantitative PCR

Extraction of RNA was achieved using a peqGOLD MicroSpin Total RNA Kit (Peqlab), which was then converted into cDNA with a High-Capacity cDNA Reverse Transcription Kit (Thermo). qPCR was performed using DyNAmo ColorFlash SYBR Green qPCR Master Mix (Thermo) and a LightCycler® 480 Instrument II (Roche). Signals of amplified products were verified using melting curve analysis and mRNA levels were normalized to Beta-Actin. Relative expression levels were calculated using the ΔΔCt method described elsewhere [36]. Primer sequences used for analyzing mRNA content were: CamKIIδ (PrimerBank ID: 26333029a1), forward ‘5-CTAGGGACCATCAGAAACTGGA -3’ and reverse ‘5-GGATCTGCTGAATGCAATGACTG -3’.

### Mouse models

Wild-type C57BL/6, male mice were purchased from Charles River Laboratories (Wilmington, MA, USA) and streptozotocin (STZ) treatment was performed as previously described [37]. Age-matched, untreated mice served as controls. Blood glucose was adjusted with insulin glargin (Lantus®, Sanofi) to <350 mg/dl on a weekly basis. Male db/db mice (C57BL/6N-Leprdb) and respective controls (db/m) were also purchased from Charles River Laboratories (Wilmington, MA, USA). All mice received water and food ad libitum. Mice were sacrificed using carbon dioxide, perfused with 0.9 % sodium chloride, and the organs immediately isolated for analysis. All procedures were approved by the Animal Care and Use Committee at the regional authority in Karlsruhe, Germany (G319/14 and G295/15). Generation of CaMKIIδ KO mice was described previously [38]. Animals received a standard diet and were maintained on a 12h light and dark cycle at a room temperature of 22 ± 2 °C and room humidity of 55%. All experimental procedures were reviewed and approved by the Institutional Animal Care and Use Committee at the regional authority in Karlsruhe, Germany (35-9185.81/G-7/15).

### Statistical analysis

Statistical data analysis was performed using GraphPad Prism 7 (GraphPad Software Inc.). All data are expressed as mean values ± standard deviation and were analyzed for significance using two-tailed unpaired t-test with Welch’s correction. The comparison of more than one group was achieved using an ordinary one-way or two-way ANOVA analysis followed by comparing all groups using Tukey’s (one-way ANOVA) or Sidak’s (two-way ANOVA) multiple comparison test. Differences were considered significant at p < 0.05. For all kinetic analyses, the data were fitted by nonlinear regression using the GraphPad PRISM 6 software (GraphPad Software Inc.), and *Km* and *V*max values were calculated.

## Acknowledgements

This study was supported by the *Deutsche Forschungsgemeinschaft* (DFG; SFB1118) and the *Deutsche Zentrum für Diabetesforschung* (DZD).

## Author Contributions

J.M., S.K., T.F., A.T., J.B. and PN designed experiments. J.M., J.K.H., J.C., S.K., performed experiments and collected the data. J.Z., M.C.C., A.S. and F.G.C. analyzed the data. J.M., T.F., A.M., A.T., P.N. and J.B. conceived and discussed the strategy about ongoing experiments. J.M., T.F. and P.N. wrote the manuscript, which was edited by all co-authors.

---

**Supplementary Figure 1.**
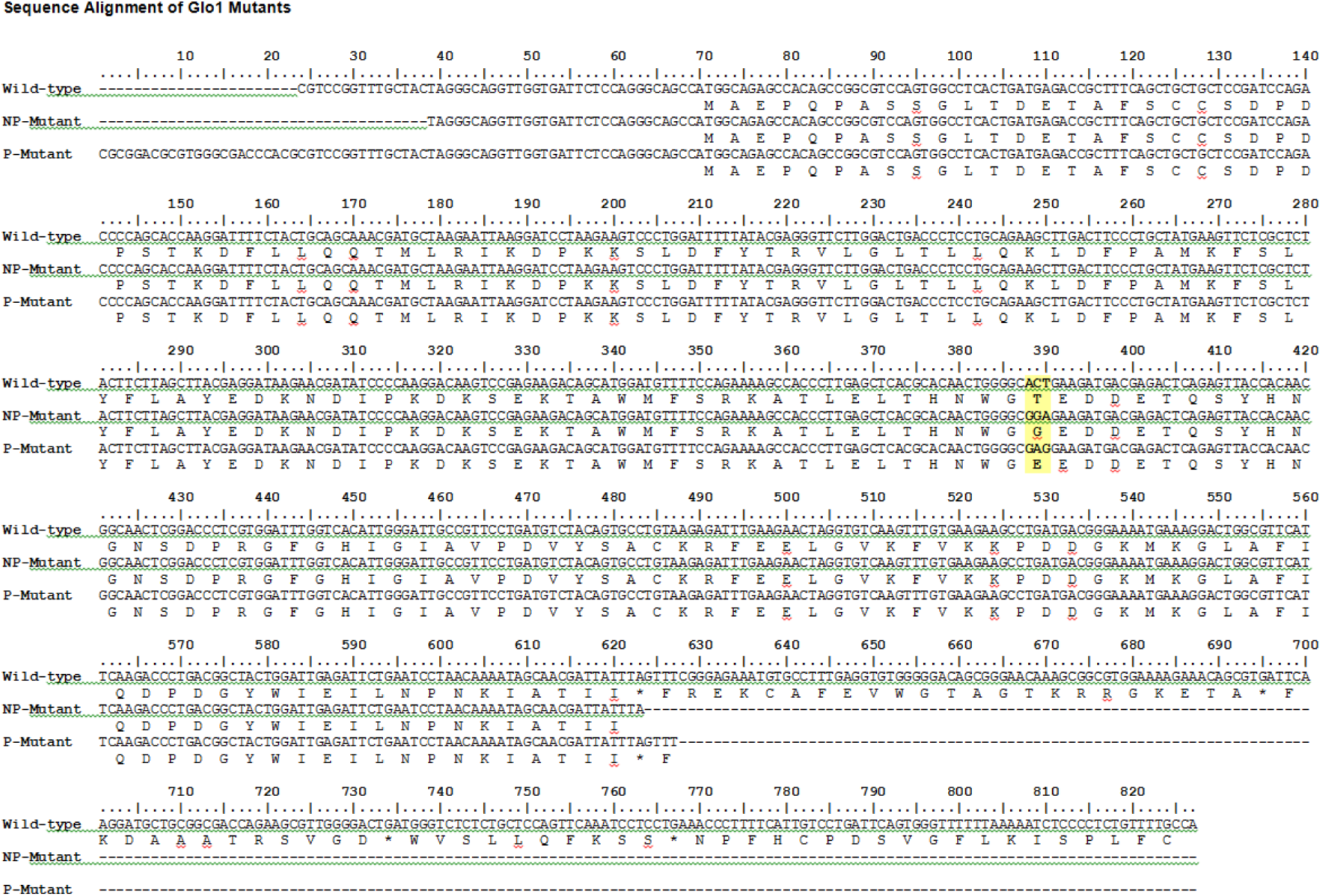
Sequence Alignment of Glo1 mutants

